# Does the heart forget? Modulation of cardiac activity induced by inhibitory control over emotional memories

**DOI:** 10.1101/376954

**Authors:** Nicolas Legrand, Olivier Etard, Anaïs Vandevelde, Mélissa Pierre, Fausto Viader, Patrice Clochon, Franck Doidy, Denis Peschanski, Francis Eustache, Pierre Gagnepain

## Abstract

Effort to suppress past experiences from conscious awareness can lead to forgetting. It remains largely unknown whether emotions, including their physiological causes, are also impacted by such memory suppression. In two studies, we measured in healthy participants the aftereffect of suppressing negative memories on cardiac response. Results of Study 1 revealed that an efficient control of memories was associated with a long-term inhibition of the cardiac deceleration normally induced by disgusting stimuli. Attempts to suppress sad memories, on the opposite, aggravated cardiac response, an effect that was largely related to the inability to forget this specific material. In Study 2, we found using electroencephalography that a prominent neural marker of inhibitory control, a suppression of the 5-9 Hz frequency band, was related to the subsequent inhibition of the cardiac response. These results demonstrate that suppressing memories also influence the cardiac system, opening new avenues for treating intrusive memories.

## 1 Introduction

Intrusive memories often take the form of distressing images and physical reactions that interrupt ongoing mentation (Brewin et al., 2010). A difficulty to control those inappropriate distressing emotional responses, which may follow or initiate intrusive mental images, is a central concern for most psychiatric conditions (Gross, 2015). Various symptoms of anxiety-related disorders comprising worries, the recall of traumatic experiences in PTSD (Shepherd and Wild, 2014) or unwanted thoughts in obsessive-compulsive disorders (Fergus and Bardeen, 2014), as well as ruminations observed in depression (Joormann and Gotlib, 2010) are representative of such difficulties. Due to historical presumptions about the independent and pernicious persistence of suppressed memories, it is often assumed that attempt to exclude from awareness those distressing images is harmful (Dunn et al., 2009). By this view, suppressed memories may backfire, causing psychological symptoms (Moritz et al., 2014), or possibly aggravating autonomic responses that accentuate perceived distress (Campbell-Sills et al., 2006). Alternatively, the origins of excessive autonomic and distressing reaction linked with intrusive memories may arise not because suppression is intrinsically bad, but because the cognitive computations achieved by the memory control network to deploy the necessary functional coping skills are disrupted or inappropriate. Here, we examine whether retrieval suppression of unpleasant images might also influence a prominent marker of emotional response: the heart rate activity.

Although targeting different components, the control of intrusions and emotions is believed to be underpinned by common neurocognitive processes orchestrated by the prefrontal cortex (Depue et al., 2015), acting protectively like an “*immune system of the mind”* (Cole et al., 2014). In addition to inhibitory modulation of areas supporting memory retrieval such as the hippocampus (Anderson, 2004; Depue et al., 2007) or the visual cortex (Gagnepain et al., 2014), functional magnetic resonance imaging (fMRI) studies have shown that suppressing negative memories is associated with down-regulation of the emotional system, such as the amygdala (Depue et al., 2010; Gagnepain et al., 2017; Depue et al., 2007). (Gagnepain et al., 2017) further showed that the better people were at suppressing intrusions, the more it reduced their perceived emotional states to suppressed images. These dual effects on memory and emotion originated from a common right prefrontal cortical mechanism that down-regulated the hippocampus and amygdala in parallel. Critically, a similar top-down modulation originating from the pre-frontal cortex is also seen during the direct regulation of emotional response to negative stimuli (Hayes et al., 2010), with a specific target on the amygdala (Comte et al., 2016).

Together, these results suggest that inhibitory control, the ability to stop or override a specific response comprising distressing thoughts and emotional experiences, may have strong clinical relevance in the context of various psychiatric disorders. Notably, if previous works suggest a possible beneficial effect on the cognitive and experiential level of emotions, this hypothesis questions the influence of cognitive control over other central characteristics of emotion and mental distress: being a set of automated physiological reactions. In the late 19th century, the psychologist and philosopher William James referred to emotions as feelings accompanying bodily changes (James, 1884). Several kinds of researches have since supported and completed this broad framework of embodiment, positing bodily afferent signals as the roots of emotional construction (Garfinkel and Critchley, 2016; Babo-Rebelo et al., 2016). From a physiological perspective, the activation of the autonomic nervous system (ANS) and its influence on cardiac activity is one of the most prominent manifestations of emotional reactions (Kreibig, 2010). Indeed, changes in the duration between successive heartbeats, referred as the *heart rate variability* (HRV), is representative of both sympathetic and parasympathetic influences reflecting affective responses (Valenza et al., 2014).

The *neurovisceral integration model* (Thayer and Lane, 2000) proposes that cardiac vagal control is orchestrated via a regulation of the amygdala activity by higher-order brain areas in the cortical hierarchy, such as the dorsolateral (Winkelmann et al., 2016) and medial (Sakaki et al., 2016) prefrontal areas, also engaged during memory control. In this model, the amygdala is a major efferent source of modulation of cardiovascular response via the (parasympathetic) vagal nerve. The *polyvagal theory* (Porges, 1995) further proposes that efferent vagal connections mediating parasympathetic modulation of heart activity, include a phylogenetically more recent ventral branch. This ventral branch of the vagus nerve, which may be distinctly mammalian, could selectively modulate heart activity depending on cognitive context, thus integrating attention, emotion or memory inputs in support of autonomic regulations.

Giving these lines of evidence, we propose that the cognitive control network engaged to regulate the mnemonic and emotional content of unwanted memory traces through the suppression of hippocampus and amygdala activity, respectively, may also inhibit later cardiac responses to suppressed stimuli. To test this hypothesis, we measured in two consecutive studies the electrocardiographic (ECG) response to negative and neutral scenes before and after the TNT paradigm (Fig.1). During the TNT task, participants attended to a reminder (an object) of these emotional and neutral scenes which cued them to retrieve the scene (Think items), or to suppress its retrieval (No-Think items). During pre- and post-TNT emotional evaluation, participants rated the Think and No-Think scenes’ valence, along with previously studied Baseline scenes not presented during this TNT phase. A cued-recall task was also performed immediately after the TNT task to estimate the aftereffects of memory suppression on recall performances. Study 2 additionally recorded concurrent electroencephalographic (EEG) response during the TNT task. The main goal of EEG recording was to confirm that the modulation of the cardiac response following memory suppression was truly related to the neural response engaged during the retrieval suppression of memory processes. To this end, we track the well known oscillatory dynamics supporting the inhibitory control of memories and estimate their role in the inhibition of the later cardiac response. Although the beta frequency band is frequently associated with controlling and stopping motor action (Wagner et al., 2018), here we focused on the decrease of the theta frequency band which has been consistently reported during the suppression of memory awareness using inhibitory control (Waldhauser et al., 2014; Ketz et al., 2014). This decrease is believe to reflect the suppression of mnemonic activity and should therefore be related to cardiac modulation if both phenomena are indeed intertwined as proposed here.

**Figure 1:**
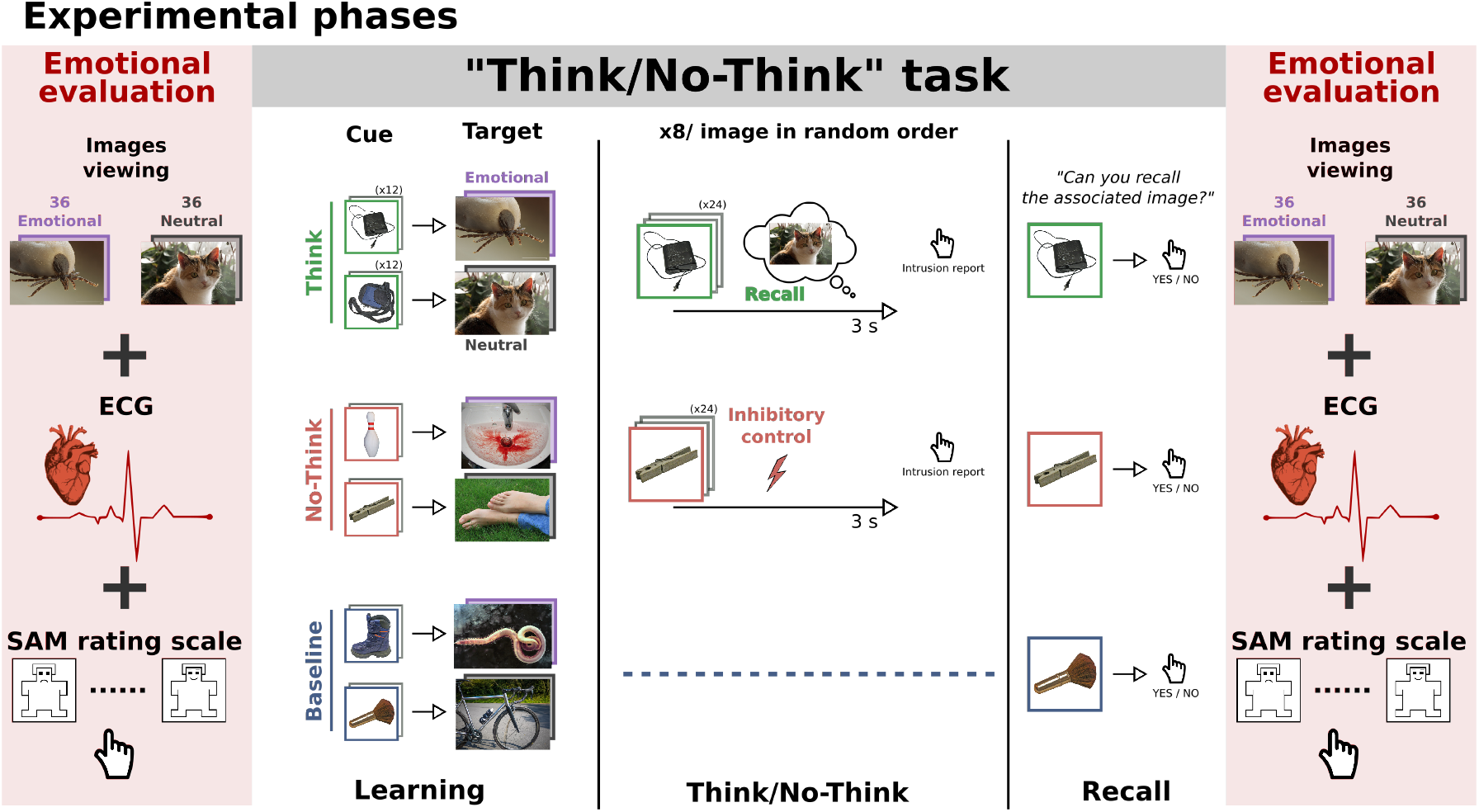
Experimental phases. After learning pairs consisting of a object and scene pictures, participants performed the think/no-think (TNT) task. The associated scene had either neutral or negative valence. For think items (in green), participants recalled the associated scene. For no-think items (in red), they tried to prevent the scene from entering awareness. The baseline items (in blue) were not presented during the TNT task. We measured subjective (i.e. valence rating) and cardiac reaction toward the scenes before and after the TNT procedure (pre-TNT and post-TNT evaluations). A recall task was also integrated after the TNT phase to estimate memory suppression. 7 details the methodological differences between Study 1 and 2.

## 2 Results

### 2.1 Results Study 1

#### 2.1.1 Behavioral results

##### Intrusions

When successful, the inhibitory control of memory recall is characterized by a progressive decrease in intrusion frequency (van Schie and Anderson, 2017). Here, we compared the frequency of intrusion of Disgust, Sadness and Neutral items over the 8 TNT blocks for the No-Think condition (Fig. 2 **A**). An Emotion x TNT blocks ANOVA showed both an effect of Emotion (F_(1.60, 43.18)_= 3.82, 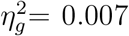, *p* = 0.04) and TNT blocks (F_(3.21, 86.62)_= 21.00,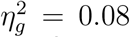, *p <* 0.001) but no interaction between these two factors (F_(7.41, 200.10)_= 0.78, 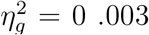, *p* = 0.62). Planned comparisons revealed that participants had on average less intrusions associated with disgusting images as compared to both neutral (t_(27)_=2.45, *p* = 0.02, *d* = 0.46, two-tailed) and sad stimuli (t_(27)_=2.04, *p* = 0.05, *d* = 0.38, two-tailed).

**Figure 2:**
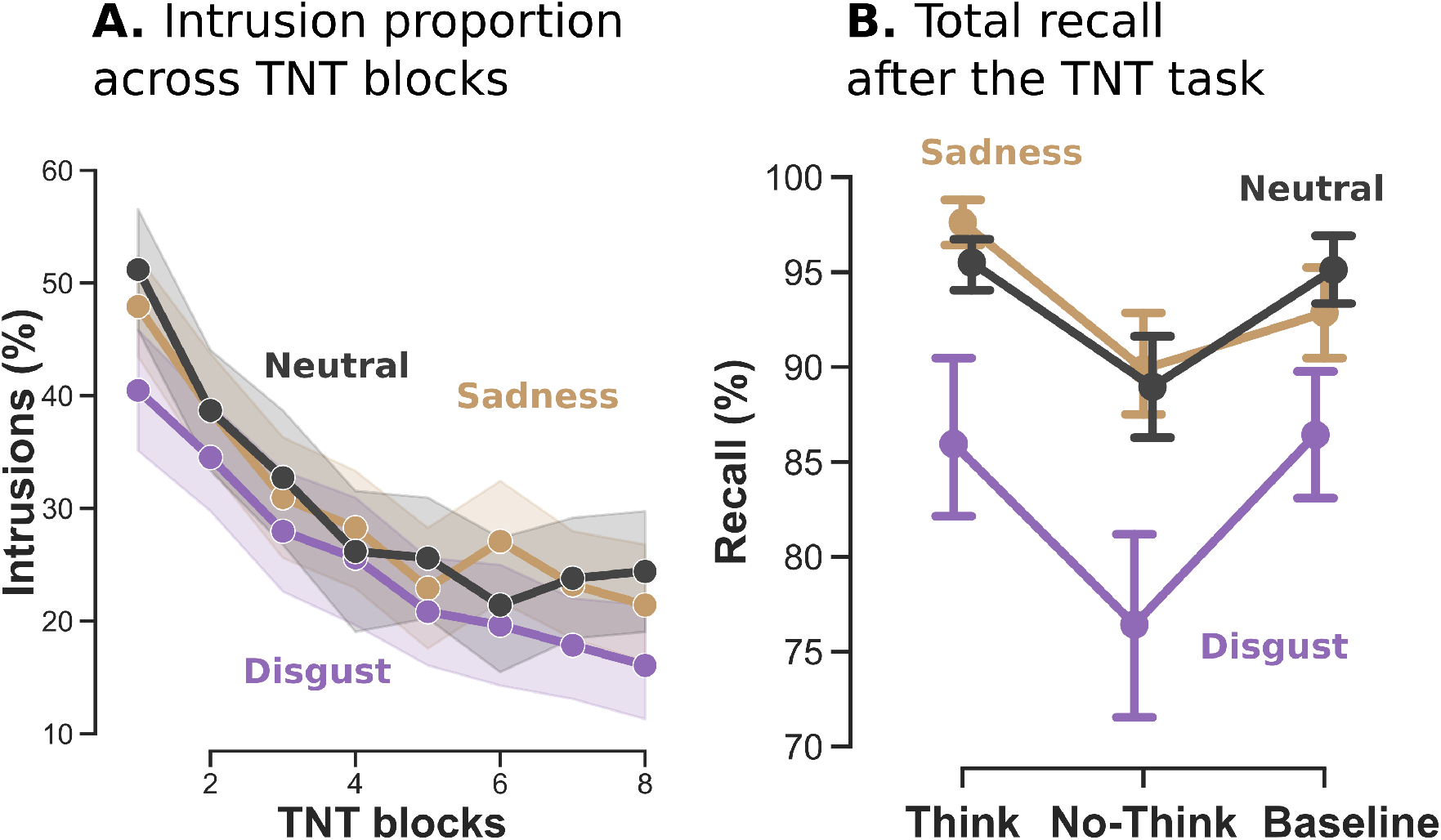
Behavioral indices of inhibitory control over memory in Study 1 (n = 28). **A.** Intrusions proportions for No-Think trials (i.e. proportion of trials where the associated memory entered into awareness while participants were instructed to inhibit recall) over the eight TNT blocks. Participants increased their ability to control intrusion over the eight TNT blocks. On average, disgusting pictures were reported less intrusive than sad or neutral ones. **B.** Total recall after the TNT procedure. Images in the suppression condition (No-Think) were more forgotten than baseline for both disgusting and neutral emotion. However, this effect was not significant for sad stimuli.

##### Inhibition of recall induces forgetting

Preventing memory from entering awareness during the TNT task is associated with a higher forgetting of the suppressed scenes during the post-TNT recall task, compared to baseline items (Anderson and Green, 2001). We first sought to replicate the standard finding an performed an Emotion× Condition [Think, No-Think, Baseline] ANOVA on recall performances (percentage of correctly recalled scenes – see Method). This analysis (Fig. 2 **B**) revealed a significant effect of both Emotion (F_(1.32, 35.51)_= 16,01, 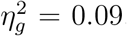, *p <* 0.001) and Condition (F_(1.74, 47.04)_= 8.17, 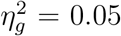, *p* = 0.001), but no interaction between those two factors (F_(2, 61, 70.44)_ = 0.86, 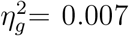, *p* = 0.45).Planned comparisons showed that participants recalled significantly less No-Think items than Baseline for Disgust (t_(27)_ = −2.27, *p* = 0.03, *d* =-0.43) and Neutral condition (t_(27)_ = −3.12, *p* = 0.004, *d* = −0.58) while no difference was found for Sadness (t_(27)_ = −0.96, *p* = 0.34, *d* = −0.18).

Taken together, these findings are inconsistent with the notion that negative stimuli may be more difficult to control and are in line with previous works reporting numerically less frequent intrusions (Gagnepain et al., 2017) and greater memory suppression (Depue et al., 2007) for negative memories. On one hand, this pattern of behavioral findings can reflect the existence of a greater motivational force driving the need to control and suppress memory for disgusting pictures. On the other hand, it may also reflect the greater natural need to avoid disgusting pictures and reduce their storage (Medina et al., 2016), which is compatible with the fact that we observed an overall poorer recall of disgusting stimuli compared to sad or neutral scenes. However, although we cannot tease part those two hypotheses in the current study, it is worth mentioning that they are not mutually exclusives and both rooted from the natural desire to avoid and reject emotional disgust.

##### Effect of suppression in the subjective evaluation

Beyond this well-documented suppression of memory representation, we also questioned whether the ability to control intrusion could relate to changes in the subjective emotional judgment. We then examined the SAM valence rating after the Think/No-Think procedure adjusted by pre-TNT evaluations (7). An Emotion × Condition ANOVA revealed no significant effect of Emotion (F_(1.70, 45.87)_ = 0.28, 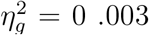, *p* = 0.72), Condition (F_(1.95, 52.57)_ = 1.17, 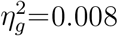, *p* = 0.32), or the interaction between those two factors (F_(3.63, 98.10)_ = 0.93, 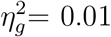, *p* = 0.44). Planned comparisons between No-Think and Baseline conditions revealed no significant difference for Disgust (t_(27)_ = 0.62, *p* = 0.53, *d* = 0.18), Sad (t_(27)_ = 0.75, *p* = 0.45, *d* = 0.14), or Neutral pictures (t_(27)_ = 0.14, p = 0.88, d = 0.02). Thus, suppression did not consistently affect the perceived valence of the scenes. An image’s perceived valence would be expected to depend, however, on the context and the valence of surrounding pictures which may modulate decision processes and affective rating. The block procedure used for Study 1, bring together items from Think, No-Think, or Baseline conditions, and decision process may therefore be confounded by the context of their occurrence. Moreover, the binary yes/no intrusion scale used in Study 1 (instead of a more subtle 3-points scale) may also have reduced the sensitivity and the awareness towards intrusions, reducing the engagement of adaptive control processes to purge momentary awareness and inhibition of the perceived valence. These potentials issues are addressed in Study 2.

#### 2.1.2 Physiological results

##### Sadness and disgust emotions decrease heart rate

We then asked if repeatedly suppressing unwanted memories across blocks of the TNT task was accompanied by persisting modulation of the cardiac response that is normally elicited by the presentation of negative scenes. Before addressing this question, we first used the pre-TNT evaluation of emotional response to characterize and isolate the specific cardiac autonomic response elicited by negative affect and unpleasant scenes. Here, we analyzed the heart rate variations evoked for each scene up to 10 seconds following the onset of the picture presentation. Such event-related analysis of cardiac response has been successfully used in previous works to characterize and quantify the heart autonomous response to emotion (Valenza et al., 2014). This method also offers the advantages to focus on the period preceding the emotional valence rating which reflects more precisely the item specific emotional content. In addition, this method also offers a greater flexibility to deal with spontaneous artifacts which are problematic in the context of a block analysis. Finally, it also reduces the influence of the respiratory signal whose contamination may be more apparent when cardiac activity is averaged over longer block duration.

We compared the averaged BPM curve across the three emotional conditions in the first pre-TNT evaluation (see Fig. 3 **A**). Results show that presenting scenes categorized as Disgusting (maximum deflection peak = −2.34 at 3.48 s, t_(27)_ = 4.57, P-FDR *<*.01, twotailed paired t-test) or Sad (maximum deflection peak = −2.24 at 3.36 s, t_(27)_ = 4.60, P-FDR *<*.01, two-tailed paired t-test) induced a significant deceleration of the heart rate compared to Neutral pictures (correction for multiple comparisons across time-points at P-FDR¡.05). This deceleration in response to negative scenes occurred within an emotional time of interest (TOI) ranging from 1.89 to 5.03 seconds.

**Figure 3:**
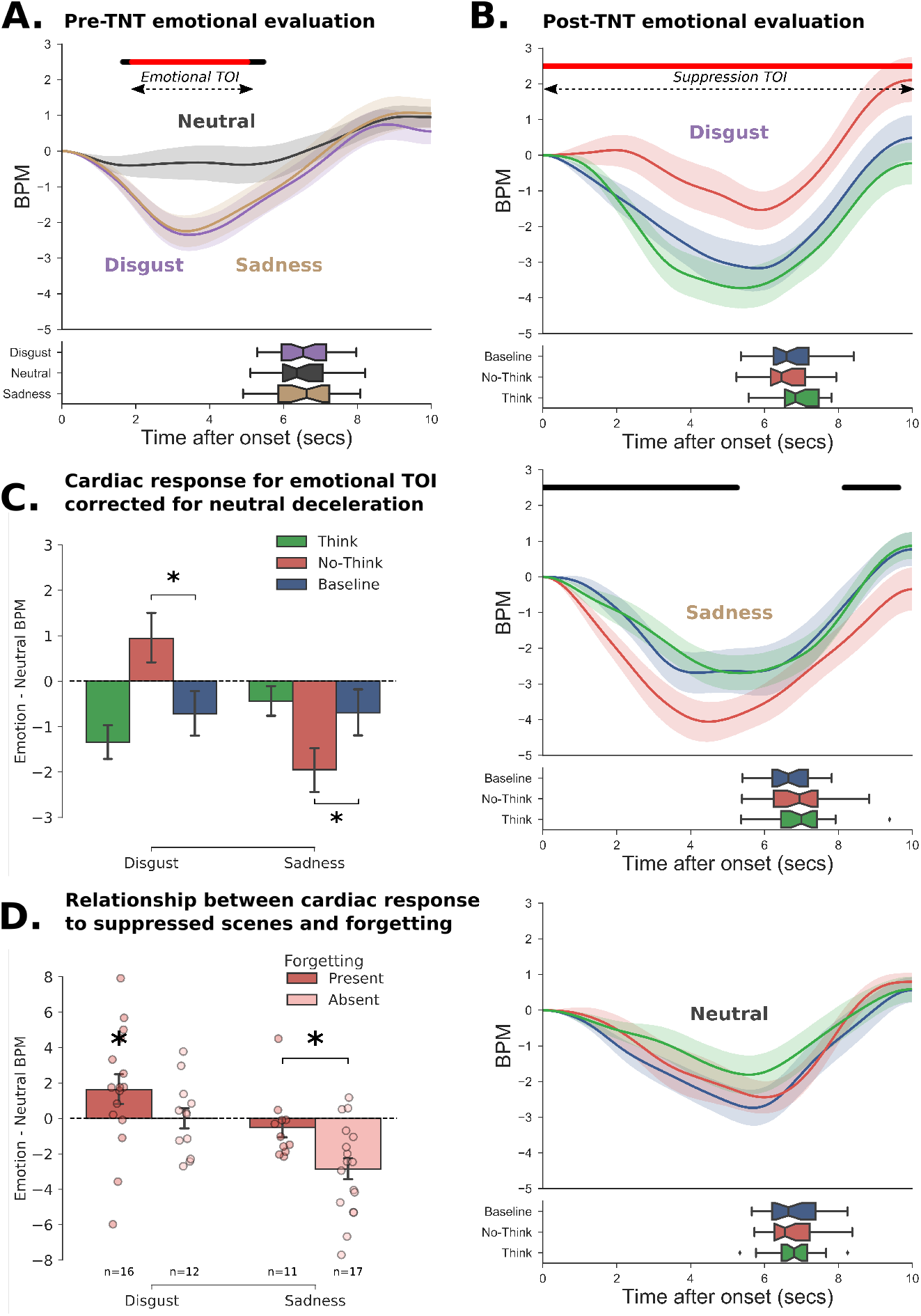
Cardiac response toward stimuli before and after the TNT procedure (Study 1). **A**. Cardiac response following the initial presentation of the images before the TNT task. The curves represent the changes in the number of heartbeat per minutes (BPM) using the onset of image presentation as a zero starting point. In the lower panel, the boxplots represent the distribution of response times for the valence rating. **B**. Cardiac response after the TNT task for Disgust (top panel), Sadness (middle panel), and Neutral (bottom panel) images. For each emotion, we represent Think, No-Think and Baseline items separately (green, red and blue lines respectively). We observed a significantly reduced deceleration of the heart rate for disgusting No-Think items compared to baseline, while a greater deceleration was reported for sad picture in the suppression condition. For both panel A and B, the black line shows the significant difference between neutral and emotional (i.e. Disgust + Sadness) evoked response, while the red line indicates the remaining significance after FDR correction. **C**. To control for the stronger deceleration observed for neutral items in the post-TNT evaluation, we computed the difference between the average heart rate of emotional and neutral pictures over the significant emotional time window extracted in the pre-TNT evaluation. The reduced or increased deceleration observed for No-Think items was still present for Disgusting and Sad pictures, respectively, even after controlling for the neutral deceleration. This suggests that the modulation of the cardiac response following memory suppression cannot be accounted by trivial attentional orientation processes. **D**. To explain the opposite modulation observed for Disgust and sadness, we split participants according to whether they could suppress or not emotional scenes from their memory. Participants who were better at suppressing Disgusting scenes from their memory also showed a reduced deceleration of cardiac response toward them afterward. On the opposite, the increased deceleration of cardiac response following the suppression of sad items was mostly related to participants who could not suppress them from their memory.

##### Suppression of unwanted memories inhibits later cardiac deceleration linked with disgusting scenes

We then questioned if the repeated cognitive control over emotional memories affected this typical heart rate deceleration that we observed in the pre-TNT evaluation, by comparing the heart rate changes in the post-TNT evaluation following the presentation of Think, No-Think and Baseline scenes for the three Emotional conditions (see Fig. 3 **B**). We predicted that suppression would reduce the cardiac response normally characterizing negative scenes. This would be reflected by less deceleration of the cardiac response following the presentation of an unpleasant scene previously suppressed and thus a positive autonomous response suppression difference compared to Baseline (No-think – Baseline time-courses). For Neutral scenes, we did not have specific predictions as to whether retrieval suppression should further remove residual negativity (which might actually increase bpm rate following the Neutral scene), or, instead, simply dampen positive affect for the item and thus perhaps shows a more pronounced deceleration. Given the results observed in the pre-TNT emotional evaluation, which shows a larger deceleration of heart rate following the presentation of negative scenes over a 1.89 to 5.03 seconds time-window, we hypothesized that the effect of memory suppression over the cardiac response should arise over a similar time-window and would induce a lesser deceleration of heart rate. This hypothesis was confirmed for Disgusting stimuli, for which we found the expected reduced deceleration compared to Baseline conditions, over the entire interval of presentation with the peak of maximum difference observed at 4.68 seconds (t_(27)_ = 2.41, P-FDR = 0.029, *d* = 0.45). This pattern of results was not observed for emotional Sadness for which No-Think items (−4.05 bpm after 4.48 seconds) augmented bpm deceleration compared to Think (−2.69 bpm after 5.33 seconds) or Baseline stimuli (−2.68 bpm after 4.16 second). However, the observed difference with baseline did not survived multiple comparisons correction (t_(27)_ = 1.98, P-FDR = 0.06, *d* = 0.37, one-tailed). Concerning the Neutral condition, paired t-test contrasting Think (−1.8 bpm after 5.57 seconds) and No-Think (−2.44 bpm after 5.99 seconds) with Baseline (−2.73 bpm after 5.68 seconds) did not show any significant difference (all Ps *>* 0.45, one-tailed).

The cardiac modulation associated with the neutral condition was clearly different in the post-TNT session compared to the pre-TNT session. Indeed, a deeper deceleration was now observed for neutral scenes which was clearly absent during the pre-TNT phase. This discrepancy may arise from a significant difference in cardiac frequency between these two measurements. The averaged heart rate over the periods of interest was much higher during the first evaluation (80.58 *±*10.86 bpm) than during the second one (70.80 *±* 9.77, (t_(27)_ = −5.93, *p <*10e-5). This difference can be explained by the prolonged seated position during the protocol as well as a higher level of stress in the first session due to the unusual experimental situation for the participants. This diminution of about 10 bpm is enough to increase global HRV, as this measure is dependent to the initial heart rate (Mccraty and Shaffer, 2015). A higher frequency and lower HRV reduce the time between successive beats, leaving less room for heartbeats to be modulated by attentional and stimuli processing (Abercrombie et al., 2008). As a consequence, individuals with higher resting HRV often show a deeper deceleration when confronted to disgusting stimuli (Ruiz-Padial et al., 2017). Here, our results are coherent with this observation as the global deceleration peak across conditions observed in the second post-TNT evaluation is more pronounced (up to −4 bpm) than in the first evaluation.

We then wanted to control for this common deceleration reflecting stimuli processing in order to isolate the specific emotional response. We subtracted the neutral deceleration (averaged across Think, No-Think and Baseline items) from the deceleration waveform of each emotional condition and then averaged this difference over the emotional time of interest identified during the pre-TNT evaluation (see Fig. 3 **C**). Critically, even after controlling for this common neutral deceleration, comparisons between TNT conditions revealed that the memory control condition (No-Think) reduced the subsequent cardiac deceleration in the targeted time-window compared to Baseline (t_(27)_ = 2.65, *p* = 0.01, *d* = 0.50), while it induced a stronger deceleration for sad stimuli, which was also significantly lower than baseline for this emotion (t_(27)_ = −2.21, *p* = 0.03, *d* = 0.41). This pattern evidenced a clear interaction between memory suppression and emotional category on the subsequent modulation of cardiac activity (t_(27)_ = 4.25, *p <* 0.001, *d* = 0.8).

Thus, while memory suppression seems to benefit to disgusting scenes and abolishes their impact on the cardiac response (i.e. less deceleration), it paradoxically increases the cardiac deceleration observed for sad scenes. Previous work has reported that the capacity to regulate memory content and affective response are intimately related. However, as we reported in Fig 3, sad images, compared to disgusting ones, were both more difficult to control (higher intrusion rate) and less forgotten following repeated suppression attempts. We therefore hypothesized that the opposite suppression effects observed between emotional categories could, in fact, reflect a difference in the ability to control those images. When unsuccessful, inhibitory control may indeed worsen the emotional response associated with the targeted item (Gagnepain et al., 2017). From this perspective, the persistence of sad pictures even after repeated attempts to suppress them would be accompanied by a stronger emotional response, while the successful control of disgusting scenes could yield a parallel inhibition of both memory and cardiac activity.

To assess this hypothesis, we split the participants into 2 groups according to whether or not they had forgotten the No-Think items following the TNT procedure. For each emotion, we selected participants who had forgotten at least one out of the six items (Forgetting present group; *n* disgust = 16, *n* sadness = 11). The second group included participants who recall correctly all items in the No-Think condition (Forgetting absent; *n* disgust = 12, *n* sadness = 17) (see Fig. 3 **D**). The ideal procedure would have required to correct the number of forgotten items with the recall of the Baseline. However, this computation led to a small group of high suppressors in the sad condition (n = 7) inappropriate for further analysis. Concerning emotional Disgust, the “Forgetting present” groups exhibited on average less cardiac deceleration for suppressed items compared to neutral deceleration (t_(15)_=1.89, *p* = 0.03, *d* = 0.47, one-tailed). This difference between the cardiac response to disgusting and neutral No-Think items was not found for the “Forgetting absent” group (t_(11)_=0.02, *p* = 0.49, *d* = 0.006, one-tailed). Concerning sad pictures, the greater deceleration was solely observed for the “Forgetting absent” while no difference reported for the “Forgetting present” group (t_(25.494)_ = −2.78, *p* = 0.01, *d* = 1.04). This pattern of findings critically suggests that the ability to suppress a mental image from memory is crucial to observe an inhibition or a reduction of the associated cardiac response. When the memory cannot be effectively suppressed as we observe for sad pictures, repeated attempts to control and exclude from awareness an unwanted memory seems, on the opposite, to have a pernicious impact and accentuate the cardiac deceleration in response to the suppressed material.

### 2.2 Results Study 2

Study 1 evidenced that cardiac response to unpleasant images can be influenced by previous repeated inhibitory control over emotional memories. Our data suggest that this effect is specific to disgust and that this may be due to greater mnemonic control abilities over this emotion compared to sadness. However, our design could nonetheless produce confounding factors explaining the modulation of cardiac activity we reported. We chose a block design for the presentation of the emotional stimuli in Study 1 (see 7). This block presentation has the disadvantage to capture low variation linked to attentional slow process even with event-related analysis. One could for example argue that, once the participant is aware to have entered the No-Think condition, his expectation toward the stimuli differs from the other conditions and thus influence ongoing cardiac activity. To control for this possible bias, we conducted a second study adopting a semi-randomized design (see 7). We kept the separation of emotional categories in time but randomized the presentation of Think, No-Think, and Baseline stimuli inside each block. Doing so, participants were only aware of the emotional valence of the forthcoming stimuli but could not predict its condition.

Moreover, to further show that the observed effect is not linked to attentional confound occurring during the emotional assessment, it is also crucial to demonstrate that the post-TNT cardiac modulation emerges as a consequence of the neural mechanism engaged during inhibitory control (i.e. TNT phase). One finding with important implications is the observation in Study 1 of substantial individual differences in the affective consequences of retrieval suppression. This observation, along with similar prior findings (Gagnepain et al., 2017), highlights the existence of inter-individual differences in using suppression as an effective regulation and coping strategy. Thus, it would be crucial to clarify whether the large inter-individual differences observed in the inhibition of the cardiac response following memory suppression relate to differences in the neural mechanisms supporting inhibitory control. We then adapted the TNT procedure utilized in Study 1 to an electroencephalography (EEG) protocol. EEG has been previously used in numerous works together with the Think/No-Think paradigm, notably focusing on the oscillatory dynamic underlying memory suppression (Ketz et al., 2014; Depue et al., 2013; Waldhauser et al., 2014). However, given that EEG requires a large number of trials per conditions, we limited Study 2 to disgusting pictures and did not include emotional sadness. This choice was further motivated by the difficulty to control and suppress this particular material as indexed by recall performances and intrusion proportion during the TNT task. Twenty-four new participants underwent this procedure. The full experimental protocol is detailed in the section **Materials and methods**.

#### 2.2.1 Physiological results

##### Study 2 confirmed the influence of memory suppression on cardiac response

Behavioral findings from Study 1 were largely replicated in Study 2 (see supplementary information for details). The only exception was now the presence of a significant suppression of the negative affect associated with the processing of suppressed negative scenes post-TNT (see A.3), suggesting that the block presentation of Study 1 may have compromised the detection of this behavioral effect. Replicating findings observed in Study 1, results show that presenting scenes categorized as Disgusting induced a significantly greater deceleration of the heart rate compared to Neutral pictures over a window of significance ranging from 0.81 to 4.64 seconds, with the peak of maximum difference observed at 3.28 seconds (t_(23)_ = 3.52, P-FDR = 0.009, *d* = 0.71, two-tailed paired t-test, see Fig. 4 **A**).

**Figure 4:**
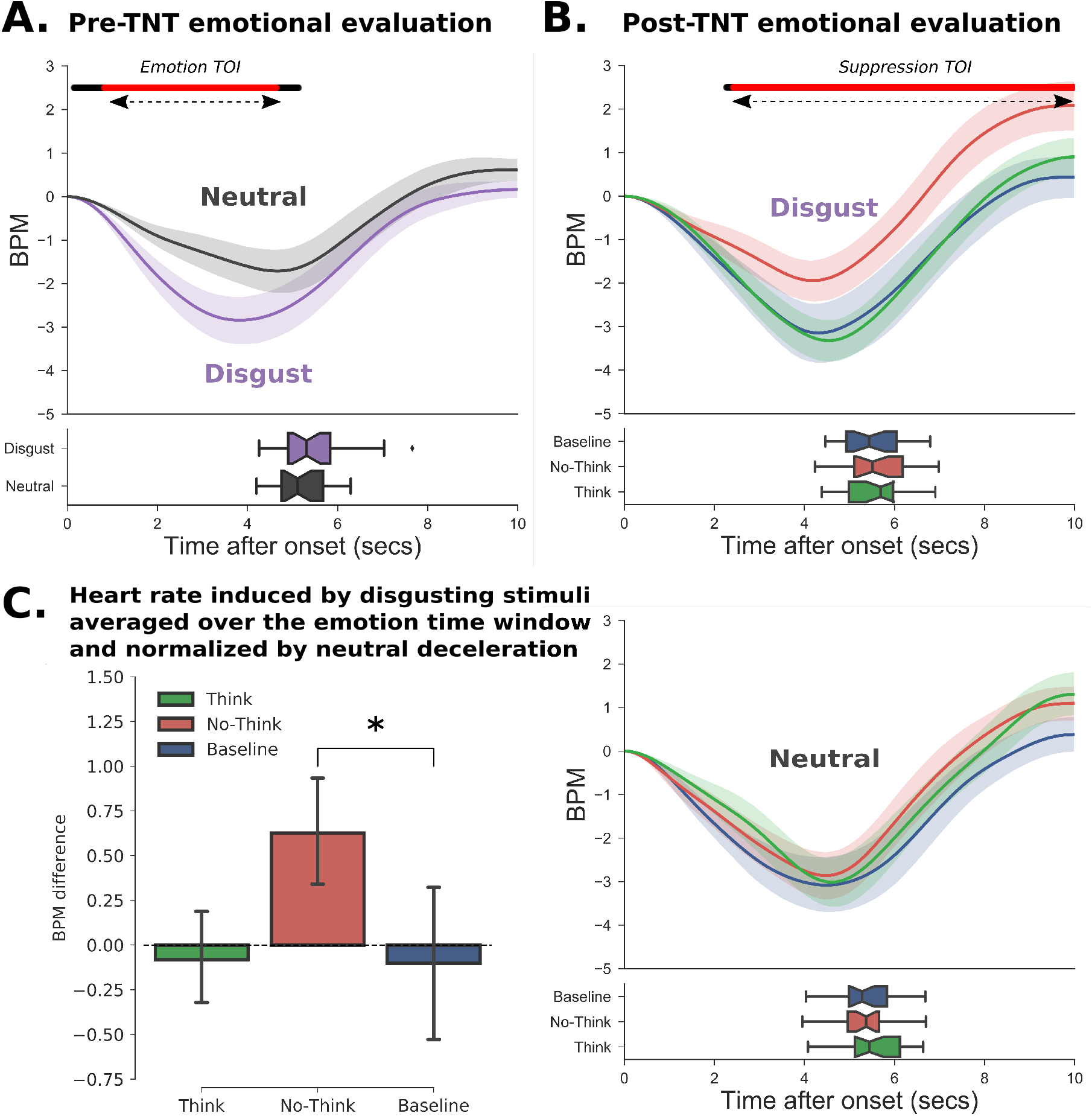
Cardiac response toward stimuli before and after the TNT procedure (Study 2). **A.** Cardiac response following the initial presentation of the images before the TNT task. The curves represent the change in the number of heartbeat per minutes (BPM) taking the onset of image presentation as a zeros starting-point. In the lower panel, the boxplots represent the distribution of response times for the valence rating. **B.** Cardiac response after the TNT task for Disgust (top panel) and Neutral (bottom panel) images. For each emotion, we represent Think, No-Think and Baseline items separately (green, red and blue lines respectively). We observed a significantly reduced deceleration of the heart rate for Disgusting No-Think items compared to Baseline. For both panel A and B, the black line shows the significant difference between neutral and disgust evoked response, while the red line indicates the remaining significance after FDR correction. **C.** The inhibition of cardiac response following memory was still observed even after controlling for the stronger deceleration observed for neutral items in the post-TNT evaluation.

We then compared the heart rate changes in the post-TNT evaluation following the presentation of Think, No-Think and Baseline scenes for the items categorized as disgusting (see Fig. 4 **B**). When comparing No-Think and Baseline conditions at each time point using FDR correction for multiple comparisons, we observed a window of significance ranging from 2.41 seconds to the end of picture presentation with the peak of maximum difference observed at 7.80 seconds (t_(23)_ = 2.45, P-FDR = 0.03, *d* = 0.50). No differences were found between conditions for Neutral scenes (all Ps *>* 0.2).

Similarly to Study 1, we observed an overall larger deceleration of the cardiac response to neutral images during the post-TNT evaluation compared to the pre-TNT evaluation. Again, this effect might be due to an attentional orientation effect on the cardiac response, which is more likely to occur given the lower cardiac frequency of the participants during the final post-TNT evaluation (bpm = 68.64 *±* 10.16) compared to the initial assessment (bpm = 78.40 *±* 8.72). Critically, the impact of memory suppression on the cardiac deceleration was still significantly different from Baseline even after controlling for neutral deceleration in the emotional time-window of interest (t_(23)_ = 2.24, *p* = 0.03, *d* = 0.45, see Fig. 4 **C**)), replicating findings from Study 1.

##### Reduction of the 5-9 Hz frequency power during cognitive control predicts cardiac modulation

We then wanted to corroborate that the observed cardiac modulation was linked to neurophysiological markers of cognitive control. We recorded the EEG signal of participants in Study 2 while they performed the TNT task. Our goal here was to quantify the inter-individual differences in controlling and suppressing the targeted memory during the TNT task. By this view, oscillatory dynamics supporting the control and suppression of memory replay should be related to later cardiac reaction inhibition. Based on similar paradigms (Ketz et al., 2014; Depue et al., 2013; Waldhauser et al., 2014), we restricted our analysis to the 5-9 Hz frequency band, whose oscillatory dynamics most likely reflects a desynchronization the medial temporal lobe. While theta activity is preferentially measured between 3 and 8 Hz, our goal here was to ground our analysis on previous findings to increase the reliability of subsequent comparisons and interpretations. The power of this oscillatory dynamic observed during the No-Think trials, contrasted with the values associated with memory recall (i.e. Think trials), would then reflect the neural suppression effect for each participant. We then related this marker to the amount of cardiac inhibition observed during the subsequent post-TNT emotional evaluation.

After extracting frequency power for Think and No-Think condition of emotional images, we contrasted these conditions in order to observe the specific decrease associated with inhibitory control using non-parametric clustering statistics (Maris and Oostenveld, 2007) as implemented in MNE python. This procedure revealed 14 significant clusters among which 1 passed the permutation statistic. This cluster (*p <* 0.001) lasted from 460 ms to the end of the trials and revealed a broad power decrease throughout No-Think as compared to Think trials (see Fig. 5 **A.**).

**Figure 5:**
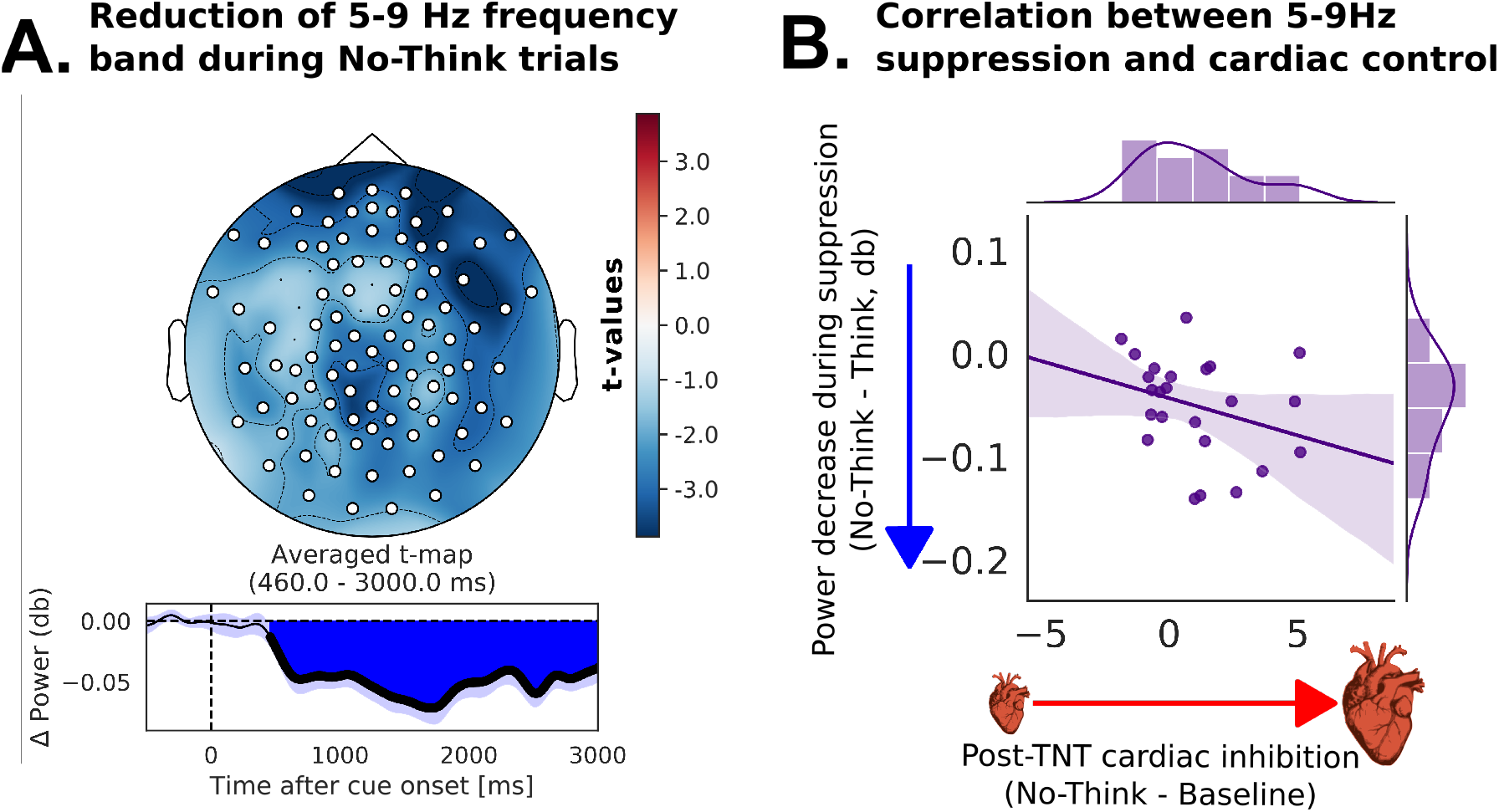
Reduction of 5-9 Hz frequency power during mnemonic control and its relationship with subsequent cardiac inhibition. **A.** Decrease of 5-9 Hz frequency power during No-Think versus Think trials. Shaded areas represent bootstrapped standard error of the mean. Only the significant electrodes are displayed on the left panel. On the right panel, the significant time-window is represented by the bold line and the blue colored area. **B.** Reduction of 5-9 Hz frequency power correlates with subsequent cardiac inhibition of disgusting stimuli. This reduction is measured as the averaged difference of power between Think and No-Think trial over the significant time-window and cluster of electrodes. The cardiac inhibition is measured as the averaged heart rate difference between No-Think and baseline items over the intersection between the emotional and the suppression time-window extracted in the pre- and post-TNT evaluation, respectively.

**Figure 6:**
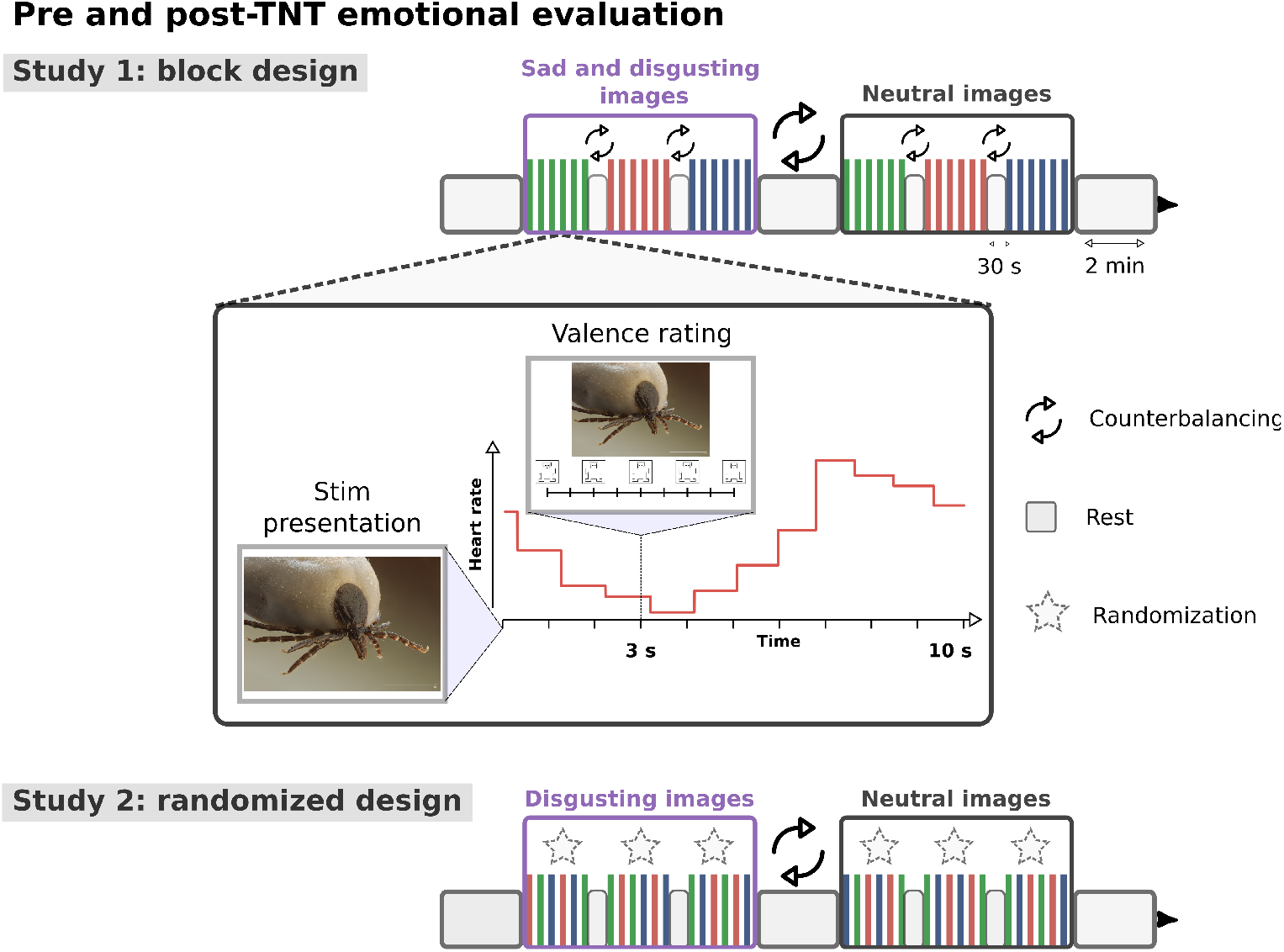
Emotional assessment in Study 1 included a block design during which stimulus presentation alternate between discrete epochs of scenes drawn from the same condition (i.e.Think, No-Think, and Baseline). This design was intended to optimize the detection of differences in heartbeats which may have only arisen in the low-frequency range. One potential problem with blocked designs, however, is that the response to events within a block may be confounded by the context of their occurrence (e.g. when participants become aware of the blocking and may alter their strategies/attention as a consequence). To control for that potential bias, Study 2 used a design in which the presentation of Think, No-Think, and Baseline items was randomized and unpredictable. However, we kept emotion and neutral conditions presented in separated blocks to control for long-term autonomic change induced by emotions that may spread over and modulate cardiac response to neutral scenes. In addition, in Study 1, an arousal scale was also presented at the bottom of the screen, ranging from 1 (corresponding to a calm face on the far left of the scale) if a picture made them feel completely relaxed, to 9 (corresponding to an excited face on the far right if a picture made them feel completely aroused). If participants felt neutral, neither relaxed nor aroused, they were then instructed to press the square under the figure in the middle position. This last change explains the difference in mean response time between Study 2 and Study 1.

**Figure 7:**
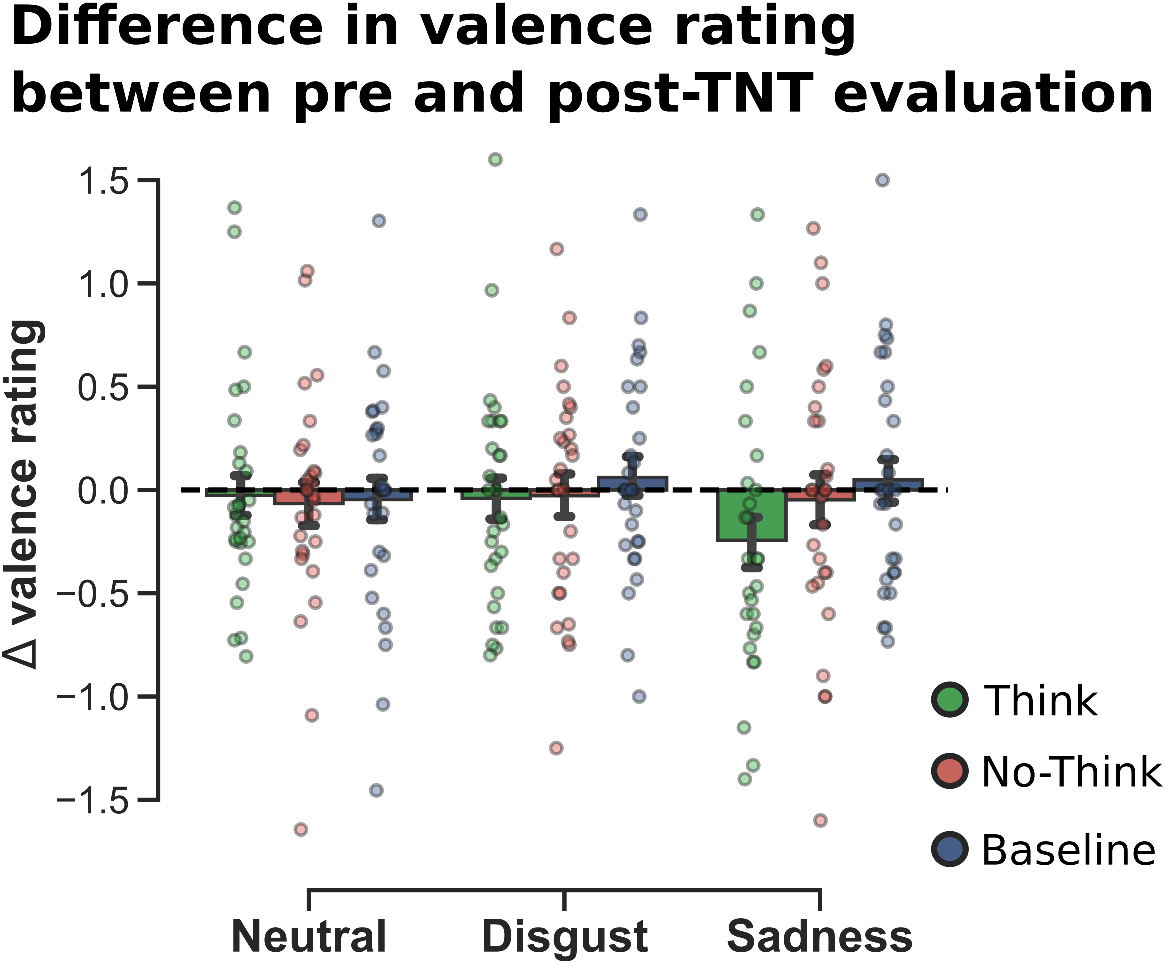
Difference in valence rating between pre- and post-TNT evaluation (Study 1). We found no significant differences in term of subjective rating for the No-Think pictures as compared to baseline in any of the three emotional conditions. Shaded error bands and error bars represent the bootstrapped standard-error of the mean.

**Figure 8:**
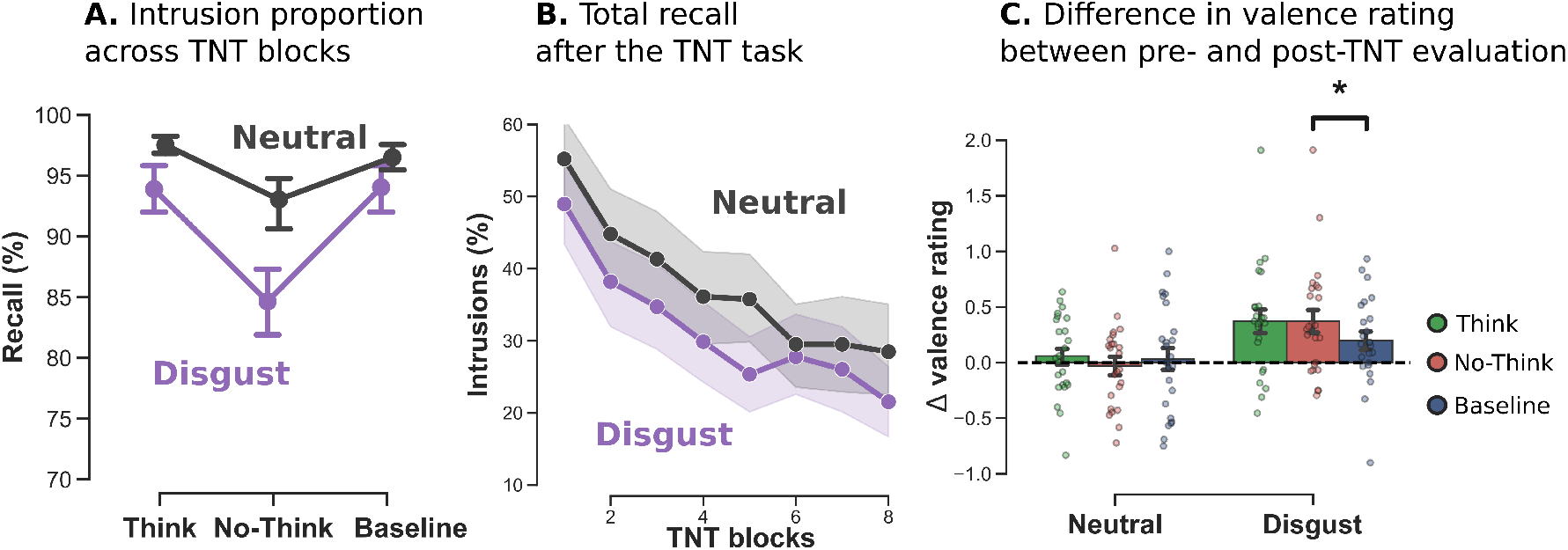
**A.** Intrusions proportions for No-Think trials (i.e. proportion of trials where the associated memory entered into awareness while participants were instructed to inhibit recall) over the eight TNT blocks. Participant increased their ability to control intrusion over the eight TNT blocks. On average, disgusting pictures were reported less intrusive than neutral ones. **B.** Total recall after the TNT procedure. Images in the suppression condition (No-Think) were more forgotten than baseline for disgusting items, an effect that was not found for the neutral emotion. **C.** Difference in valence rating between pre- and post-TNT evaluation. We found a significant difference in valence rating between No-Think and Baseline pictures for disgust, but this effect was not significant for neutral items. Shaded error bands and error bars represent the bootstrapped standard-error of the mean.

We then questioned if the suppression of frequency power during No-Think trials (compared to Think condition) was related to the cardiac modulation measured during the post-TNT evaluation. We performed one-tailed spearman’s correlation using the robust correlation toolbox (Pernet et al., 2013). We used skipped correlations to control for bivariate outliers. We found that the cardiac inhibition (i.e. lesser deceleration of the cardiac response to No-Think compared to Baseline scenes over the emotional time of interest identified during the pre-TNT evaluation) was negatively correlated (robust Spearman *r* = −0.40, *p <* 0.05, bootstrapped CI = [-0.70 to −0.01], no outliers detected) with the mean power difference between No-Think and Think trials (see Fig. 5 panel **B**). The foregoing results support the hypothesis that the neural mechanism engaged to suppress retrieval does not simply reduce the accessibility of episodic memories but can also alter the cardiac response associated with the suppressed negative memories. Individuals who adaptively suppressed the reactivation of unwanted memory traces could also disrupt emotional content of suppressed traces at the same time, decreasing their later negative impact on the physiological response.

## 3 Discussion

Intrusive and unwelcomed mental images are preponderant in numerous psychiatric disorders and can be a source of both physiological and psychological distress. The role of inhibitory control to restrain their cognitive and physiological manifestations has been debated and putatively described as worsening the emotional response or simply being ineffective. In these two studies, we showed that the successful inhibition of unwanted memories is accompanied by a long-term modulation of the cardiac activity associated with the presentation of the suppressed distressing material. Findings from Study 1 showed significant inhibition of the cardiac response toward disgusting stimuli following mnemonic control. A worsening effect was seen, however, over the processing of sad scenes. This discrepancy was explained by greater difficulties to control sad pictures and suppress them from memory, thus supporting the relationship between forgetting and heart rate modulation. To confirm the relationship between the neural mechanisms supporting the inhibitory control of memory awareness and subsequent cardiac inhibition, and control for potential design confound, we conducted a second study using only disgusting stimuli and during which the EEG signal was recorded to track down oscillatory dynamics supporting the inhibitory control of memories. An additional motivation to limit Study 2 to disgusting stimuli, was also to increase the number of trials per conditions and the signal-to-noise ratio. Replicating findings from Study 2, we demonstrated significant inhibition of cardiac activity that was specific to emotion and not observed with neutral items. Furthermore, we found a correlation between the suppression of 5-9 Hz frequency band, a prominent neural marker of memory reactivation, and the subsequent inhibition of cardiac response, thus supporting the notion that memory control and emotional regulation is supported by a common framework.

This pattern of findings consolidates previous works supporting the efficiency of direct suppression of unwanted memories. However, it extends previous works by showing that memory suppression may also impact the physiological dimension of emotions. This impact might be positive when memory control is successful, as with disgusting stimuli, but also negative if memory control is impossible, as we observe with sad memories. Our results also provide a consistent argument in favor of a top-down influence of higher level of neurocognitive hierarchies on the autonomic nervous system. We interpret our findings from the perspective of the neurovisceral integration model (Thayer and Lane, 2000) and discuss their implications for psychiatric disorders.

### 3.1 Cognitive control of distressing memories induces a long-term cardiac modulation

Our results show that the cardiac reaction toward disgusting memories is significantly reduced for suppressed items following the TNT task compared to baseline scenes. Alongside this effect on cardiac response, behavioral indices of both Study 1 and Study 2 show that disgusting images were successfully controlled during the TNT phase (being less intrusive than neutral or sad ones), and were subsequently more forgotten compared to baseline items (see Fig. 2 and A.3). This pattern of findings suggests an influence of the memory control mechanisms over subsequent cardiac responses linked to material excluded from awareness. Study 1 used a block presentation of the Think, No-Think, and Baseline pictures during the emotional evaluation which may have potentially lead to an attentional bias for No-think items producing such an effect. In Study 2, we controlled for this bias by presenting Think, No-Think, and Baseline condition using a random order. The difference in the cardiac response to suppressed and Baseline items was still observed following TNT, suggesting that this effect is tightly coupled to the actual image presented on the screen and not due to slow variation of attentional or physiological states. These findings corroborate recent studies that reported the existence of a link between the variability of the heart rate at rest and inhibitory control performances (Ottaviani et al., 2018). Such relationship has notably been observed during the “Think/No-Think task” (Gillie et al., 2013), thus supporting the notion that autonomic and cognitive controls are underpinned by a common framework. Critically, however, the current study demonstrate the control of cognitive representations associated with memory processing can subsequently lead to a long-term alteration of the cardiac reaction toward the suppressed items, going further than global autonomic traits observed at rest.

#### Cardiac inhibition is linked to successful suppression of unwanted emotional memories

Despite these evidences, results from Study 1 also show a contrasted pattern: while cardiac inhibition was found for disgusting items, suppression of sad memories induced on the opposite a stronger deceleration. Given that we observe a stronger cardiac deceleration in response to negative scenes in the pre-TNT evaluation, this post-TNT deceleration would suggest a stronger emotional reaction which follows memory suppression. Indeed, images categorized as sad were reported as more intrusive than disgusting ones and were not more forgotten than baseline following the TNT phase (see Fig. 2). This pattern of findings suggests that direct suppression was ineffective for these stimuli. This difference cannot be explained by a lower emotional valence, based on the rating provided by either the NAPS database (Riegel et al., 2015) or by our participants. Overall, sad pictures represent complex scenes strongly attached to social frameworks and pre-existing schema, which may explain the difficulty to suppress their content from memory. Previous works proposed that when unsuccessful, memory suppression could indeed worsen the distressing symptoms (Shipherd and Beck, 2005). To test the hypothesis that the greater deceleration we observed with sad items was due to ineffective memory suppression, we compared the cardiac modulation for sad and disgusting images of participants who effectively forgot at least one image in the post-TNT recall, with those who did not forget any (see Fig. 5). Results show that the amplitude of the cardiac modulation depended on the forgetting rate of the participants, suggesting that the worsening of the cardiac response occurred after unsuccessful control of sad memories. Thus, although cardiac modulation is influenced by memory suppression, the direction of this influence depends on the ability of the participant to successfully control and suppress distressing images from memory.

#### Suppression of 5-9 Hz frequency power correlates with subsequent cardiac inhibition

The current data support a tight connection between the successful control of memories and the subsequent inhibition of the cardiac response to the presentation of the suppressed scenes. However, the cardiac inhibition effect observed during the post-TNT evaluation does not necessarily imply that the memory control network has reduced some aspect of memory activation in service of affect regulation. Cardiac inhibition may instead arise, for instance, from other sources of attentional modulation engaged during the sensory processing of the suppressed scenes, but independent from the neural mechanisms engaged during repeated attempts to suppress the emotional memories. Contrary to these possibilities, however, we show that the cardiac modulation correlates with the reduction of the 5-9 Hz frequency band for trials where participants controlled their memory (NoThink) as compared to the condition of recall (Think), replicating earlier findings using the TNT paradigm (Waldhauser et al., 2014). Because this oscillatory dynamic was measured with EEG in the scalp of the participants, interpreting the contribution of different cortical sources may be done cautiously. Several works have shown that memory encoding and retrieval is supported by an increased synchronization of slow oscillatory activity (3-8 Hz), notably in the medial temporal lobe (Clouter et al., 2017). Alternatively, an increase of such activity in the frontal medial electrodes is also a strong correlate of the higher recruitment of cognitive control mechanisms (Cavanagh and Frank, 2014), that may also occur during the TNT procedure. The decrease we report here between 5 and 9 Hz might indeed reflect a mixed process of both higher neocortical inhibitory engagement and a decrease of memory replay.

Accurately disentangling these two components goes beyond the scope of this paper. However, we can notice that intrusions and their reactive intrusion control mechanisms are thought to be less frequent and less sustained during the No-Think trials than the synchronization initiated by memory retrieval during the Think trials. This is notably evidenced by the decrease of involuntary memory intrusions across the TNT blocks. Here, we contrasted the 5-9 Hz activity between Think and No-Think trials, which might reflect the interruption and desynchronization of memory activity during No-Think trials as compared to the synchronization induced by voluntary recall during Think trials, while inhibitory control oscillatory mechanisms might remain confounded by a broader low-frequency decrease. This result parallels previous findings obtained with similar TNT procedures. While (Depue et al., 2013) have reported an increase of theta power for No-Think trials, (Waldhauser et al., 2014) and (Ketz et al., 2014) provided divergent pieces of evidence showing a suppression of theta activity. This incongruence is thought to reflect the existence of an early and brief increase in proactive prefrontal cognitive control, and a more sustained reactive and adaptive suppression of theta oscillatory dynamics in the medial temporal lobe during the suppression trials (Waldhauser et al., 2014). The former effect is then presupposed to occur in the early phase of the stimuli especially when the participant can anticipate up-coming reminding cue (*<* 1 s), while the latter is observed all along the presentation of the reminding cue.

#### Implications for psychophysiology

Heart rate variability is a central manifestation of emotional experience (Kreibig, 2010). Frequency changes reflect an adaptation of cardiac activity to new environmental outcomes, which covertly influences perception and decision making (Allen et al., 2016). According to the neurovisceral integration model (Thayer and Lane, 2000), the amygdala, under the control of higher prefrontal areas (Sakaki et al., 2016), plays a key role in such process (Yang et al., 2007). This structure can trigger the acceleration or deceleration of the autonomic response via direct connections with hypothalamus and brainstem circuits and a complex interplay of connections with sympathetic and parasympathetic networks (Roelofs, 2017). For example, negative experiences, which may induce hyper-activation of the amygdala, are traditionally thought to induce heart rate accelerations. This assumption hold for fear, stress or anxiety, which require greater sympathetic recruitment to support defensive behaviors. However, several others emotions comprising disgust or sadness, and which may also activate the amygdala, are dominated by greater parasympathetic activity, inducing a cardiac deceleration (Rohrmann et al., 2009). We observed this decrease in the pre-TNT evaluation and interpreted it as a marker of emotional response toward sad and disgusting elements. Reducing cardiac activity is indeed an adaptive behavior toward various levels of threats. When facing blood-related or food-related disgusting items, this reaction could support preservation from blood loss or control of digestive reflex to prevent the ingestion of contaminated material. This is also observed with the freezing responses in many species. These reflexes are characterized by both increased sensory processing to prepare future behaviors and motor inhibition to avoid detection by predators. According to the polyvagal theory (Porges, 1995), such reactions are mediated by the parasympathetic influence of the older branch of the vagal nerve. Both Study 1 and Study 2 showed an inhibition of this decrease for No-Think items. We assume here, in line with a previous study, that the TNT task induced a down-regulation of the amygdala for the suppressed items (Gagnepain et al., 2017) that is also prone to change the associated autonomic response. The inhibition of cardiac deceleration following might thus reflect a fast modulation of parasympathetic influence coherent with the recruitment of the ventral branch of the vagus (Porges, 1995).

#### Implications for psychiatric disorders

Our results show promising evidence for physiological regulation of disgust when memory suppression via inhibitory control is effective. The emotion of disgust has a determinant role in the development and maintenance of numerous psychiatric condition like anxiety and obsessive-compulsive disorders (Olatunji et al., 2017). While this emotion does not always dominate the symptoms or induce the most salient ones, its experience can favor cognitive bias toward stimuli or situation and participates to the maintenance or increase of more dramatic manifestation like fear and state anxiety. The fear of contaminants and the experience of disgust they induce is for example central in obsessive-compulsive disorders (Brady et al., 2010). A high propensity to food-related disgust has also been described as a defensive mechanism to avoid caloric intake in anorexia nervosa (Vicario, 2013). These examples are straightforward as they refer to standard animal, blood or food-related stimuli, but it is worth pointing out that disgust also encompasses a moral dimension for humans (Vicario et al., 2017) that we did not explore here. Moreover, while depressive syndromes do not involve a high propensity of externally-oriented disgust, self-oriented disgust is considered as a hallmark of the condition (Powell et al., 2013). Although this is not clear whether more complex and resistant emotional states could benefit from training regime based on memory suppression, the current findings raise the hope that interventions focused on training mnemonic control could, in principle, reduce intrusions while dampening negative affect and physiological reaction linked to suppressed images.

By showing that retrieval suppression contributes to cardiac regulation, the present findings may offer insights into the mechanisms underlying intrusive symptoms in psychiatric disorders. Cardiac variability appears in this context as a relevant marker of both inhibitory control capabilities and emotional regulation. The regulation of cognitive representation associated with intrusive symptoms, while possibly inappropriate for resistant and complex emotional states, might however constitute a promising avenue to dampen primal emotional experiences and their physiological roots in the causal chain of psychiatric symptoms.

## 4 Methods and Materials

### 4.1 Participants

Twenty-eight right-handed native French speakers between the ages of 18 and 35 were paid to participate Study 1(14 males), and twenty-four to participate Study 2 (12 males).

They had no reported history of neurological, medical, visual, or memory disorders. The study was approved by the local Ethics Committee (CPP Nord Ouest I, N° ID RCB: 2015-A01727), and all participants gave written consent. Participants were asked not to consume psychostimulants, drugs, alcohol or coffee during the hours preceding the experimental period and to avoid intense physical activity the day before.

#### Material

The stimuli were 72 object-scene pairs plus six fillers pairs for Think/No-think practice. The objects were selected from the Bank of Standardized Stimuli (BOSS) (Brodeur et al., 2010, 2014) and depicted only nonliving and non-organic artifacts. The scenes were selected from the Nencki Affective Picture System (NAPS) database (Marchewka et al., 2013; Riegel et al., 2015; Wierzba et al., 2015a). Half of the critical scenes were normed as Neutral (36 images, mean valence: 5.79; SD valence: 0.84; mean arousal: 4.78; SD arousal: 0.71). In Study 1, the other half were normed as Negative, and induced either a feeling of Disgust (18 images, mean valence: 3.02; SD valence: 0.76; mean arousal: 6.42; SD arousal: 0.58) or Sadness (18 images, mean valence: 2.96; SD valence: 0.49; mean arousal: 6.43; SD arousal: 0.64). In Study 2 the other half contained only images inducing Disgust (mean valence: 3.15; SD valence: 0.68; mean arousal: 6.35; SD arousal: 0.48). We referred to the discrete quotation provided in the NAPS database (Wierzba et al., 2015b) to select the images. However, as we aimed at obtaining an homogeneous set where each scene depicted contents as unique as possible, we manually completed this selection with non-normed images from the NAPS. The Neutral and Negative lists were matched in terms of number of Animals, Faces, Objects, and People. Each object was carefully paired with a scene to avoid the existence of any pre-existing semantic association, which could have biased memory encoding. Three lists of 12 pairs (assigned to Think, No-Think, and Baseline conditions) were created for each valence condition and were counterbalanced so that they appeared in each TNT condition equally often, across participants.

### 4.2 Pre-TNT emotion evaluation

To measure the initial emotional reaction to selected scenes and provide a better estimation of the impact of subsequent TNT manipulation on affective responses, participants first performed an emotional rating task. This task consisted in rating both the valence and arousal associated with each visual scene using the Self-Assessment-Manikin (SAM) pictorial scale (Lang, 2008). These 9-point scales quantify participants’ self-reported valence and arousal response to each visual scene reported on a scale labeled with diagram-like manikins with varying emotional facial expressions.

Each scene was presented on grey background for a total of 10 seconds followed by a 500 ms inter-stimuli interval. During the first 3 seconds, the scene was presented without any scale and participants were instructed to carefully explore the scene and understand its meaning. Following these initial 3 seconds, the valence SAM pictorial scale appeared underneath the picture for 7 seconds (Study 1 also used an arousal scale but see 7). This scale ranges from 1 (corresponding to a frowning face on the far left of the scale) if a picture made them feel completely unhappy to 9 (corresponding to a smiley face on the far right if a picture made them feel completely happy). If participants felt neutral, neither happy nor sad or disgusted, they were then instructed to press the square under the figure in the middle. Participants selected using the computer mouse the facial expression most closely matching their perception of the valence of the scene.

We recorded the electrocardiogram (ECG) using the Biopac MP36R Data Acquisition system and its PC software AcqKnowledge 4.1 during the full duration of this task (20 min.). Although the presentation of these data goes beyond the scope of this paper, the electrodermal activity (EDA), were also recorded. All the participants undertook this emotional rating task in the morning (8 a.m) and were familiarized with the task settings using 10 independent images selected in the NAPS database. After an initial resting block of 2 min., the 72 images used in the Think/No-think procedure (including baseline items) were presented according to two blocks of Neutral and Negative scenes whose order was counterbalanced across participants. The presentation of these two main blocks was further separated by a resting block of 2 min. designed to prevent potential overlap and contamination between the cardiac responses to Negative and Neutral scenes. In Study 1, each block of Neutral and Negative scenes was further divided into three sub-blocks of 12 images (126 seconds per sub-block) corresponding to Think, No-Think, and Baseline condition and whose order was counterbalanced across participants. In Study 2, we randomized the presentation of these conditions inside the Neutral and Negative blocks (see 7). Each of these sub-blocks was followed by a resting period of 30 sec.

### 4.3 Learning and test/feedback

After this procedure, participants learned the 72 object-scenes pairs used in the Think/No-think task plus the six training pairs. Participants first learned all object-scene pairs through a test-feedback cycle procedure. After studying all pairs for 6 s each, participants were given test trials presenting the object cue for a given pair for up to 4 s and asked whether they could recall and fully visualize the paired scene. If so, three scenes then appeared (one correct and two foils were randomly taken from other pairs of the same emotional valence), and they received up to 4 s to select which scene went with the object cue. After selecting a scene or if the response window expired, a screen appeared for 1 s indicating whether the recognition judgment was correct or incorrect. In all cases (even if participants indicated that they could fully visualize the associated scene in the first step), each trial ended with the correct pairing appearing onscreen for 2.5 s. Participants were asked to use this feedback to increase their knowledge of the pair. Once all pairs had been tested, the percentage of total recall was displayed on the screen. If this percentage was inferior to 95% in at least one emotional condition, all the pairs were presented again in a randomized order. This procedure was repeated until the score was superior to 95%, thus ensuring correct encoding of the pairs and a comparable exposure to all the items. This overtraining procedure was intended to ensure that images would intrude when the cue was presented during the Think/No-think task and further reduces any encoding advantage negatively valenced scenes may have.

### 4.4 Criterion test

Following learning and after practicing with the Think/No-think task, participants performed an attentional task for 20 minutes using an entirely independent image set and aiming to address a different question (data not reported here). At this point, participants viewed a brief reminder of all of the studied pairs (3 s each), during which they were asked once again to reinforce their knowledge of the pairings. Following this pair refresher, a final criterion test was proposed where the participant had to recall the correct image in a similar way to the test/feedback procedure, but this time without feedback and only once. This allowed to isolate items forgotten before the Think/No-think task and to exclude them from the analysis.

### 4.5 Think/no-think task

Participants then performed the Think/No-think task, which was divided into 8 TNT blocks, each 5 min in length. After two consecutive blocks, a message was displayed on a black screen indicating to the participant that he/she could rest. The participant decided when he/she wanted to resume the experiment by pressing any button on the response box. Each block consisted in the presentations of the 24 “Think” and 24 “No-think” items, yielding, across the 8 blocks, 256 trials for Neutral and 256 for Negative conditions (Disgust and Sadness in Study 1, and only Disgust items in Study 2). Cues appeared for 3 s either framed in green or red, centered on a grey background. On “Think” trials, the cue was bounded by a green box, and participants were told to generate an image of the associated scene as detailed and complete as possible. On “No-think” trials, the cue was bounded by a red box, and participants were told that it was imperative to prevent the scene from coming to mind at all and that they should fixate and concentrate on the object-cue without looking away. During red-cued trials, participants were asked to block thoughts of the scene by blanking their mind and not by replacing the scene with any other thoughts or mental images. If the object image came to mind anyway, they were asked to push it out of mind.

In Study 1, after the offset of each of the Think or No-Think trial cues, participants reported if the associated scene had entered awareness by pressing the button under their right index if it did, or the button under their right middle finger if it did not. Participants had up to 2 seconds to make this rating, so they were instructed and trained to make this quickly without thinking about the associated picture. Their response was followed by a jittered fixation cross randomly lasting between 2400 to 3600 ms.

In Study 2, electroencephalographic (EEG) activity was recorded inside a Faraday’s cage during this procedure. Stimulus presentation was controlled by Eprime Pro on a 17” LCD screen with a 1280 × 1024 resolution. To avoid exploratory eye movement, each picture was displayed inside a 400 × 400 pixel square. Participants were seated comfortably in a dimly lit room during the whole experiment at a distance from 90 cm to 100 cm from the screen and were instructed to minimize blinking and moving during recording.

Participant had to give feedback concerning the intrusion rate on a response box using their right hand. The fixation cross stayed on the screen until the participant’s feedback. Participants reported the extent to which the associated scene had entered awareness by using a 3-point scale, instead of binary yes/no choice in Study 1, corresponding to the labels: never, briefly, often. Although we used participants’ responses to classify each No-Think cue as either an intrusion (i.e. a “briefly” or “frequent” response) or not (a “never” response) in a binary fashion, this 3-point scale was proposed to increase participants’ awareness toward intrusive memory and ensure that participants would engage appropriate inhibitory resources throughout the task to countermand reminiscence of memory whether the very briefly, or strongly, pops into participant’s mind. In addition, while participants had no explicit time constraint to give their feedback during this procedure, we asked them to make these reports as quickly and intuitively as possible. We do so to avoid that they dwell too much on the intrusion and finally recall the memories.

### 4.6 Recall

After this procedure, after effects of memory suppression were examined via a cued-recall task on all the object-scene pairs, including the 24 Baseline scenes (12 Neutrals and 12 Negatives) omitted from the Think/No-think task. During trials of this cued-recall task, the cue object was presented in the center of the screen for 5 seconds and participant were told to visually recall the associated picture. If they could recall the associated scene, they were told to press the button under their right index finger and then had 15s to verbally describe the scene in as much detail as possible. The descriptions were recorded. If they could not recall the associated scene, they had to press the button under their right middle finger, and the next object was presented to them. Participants’ descriptions were scored checked to ensure enough coherence between the described item and the real pictures. On average, participants made a false recall for 0.3% of the images in Study 1 and for 0.8% in Study 2.

### 4.7 Post-TNT emotion evaluation

Following the recall task, participants were asked to rate the 72 pictures using the same procedure to measure whether retrieval suppression influenced later affective responses to scenes. During this task, all of the scenes (“Think”, “No-think” and “Baseline”) were presented again and we asked participants to give their perceived emotional valance and arousal while physiological signals were recorded. This procedure enables us to determine whether retrieval or suppression altered the valence, arousal or the cardiac response associated with the item, relative to “Baseline” pairs.

### 4.8 EEG

### 4.9 Recording

Application of the EEG cap (Electrical Geodesics, INC., Hydrocel Geodesic Sensor Net) was fast (about 15mn) causing a very trivial delay between learning and TNT task. EEG activity was recorded continuously by GES 300 amplifier (Electrical Geodesics, INC.) using an EGI Hydrocel Geodesic Sensor Net (HGSN-128) with a dense array of 128 Ag/AgCl sensors (Tucker, 1993). Impedances were kept under 100 kΩ (Ferree et al., 2001), and EEG channel was referenced to a vertex reference Cz and ground to CPPZ (fixed by the EGI system). The signal was sampled at 20 kHz frequency with a 24-bit A/D and was online (hardware) amplified and low pass filtered at 4 KHz. The signal was filtered by a Butterworth low pass Finite Inpluse Response (FIR) filter at 500 Hz and sub-sampled at 1 KHz. Electro-oculogram was recorded using four electrodes placed vertically and horizontally around the eyes. EEG data were processed offline using Netstation 4.4.2 (Electrical Geodesics Inc., Eugene, OR, USA). The signal was filtered using a 1 Hz Kaiser FIR first-order high-pass filter (which ensures a linear phase and no distortion in the bandwidth) in order to discard DC and very slow waves.

### 4.10 Preprocessing

EEG analyses were performed using the version 0.15 of the MNE library (Gramfort, 2013; Gramfort et al., 2014) implemented in python 3.6 and followed recently recommended analysis steps (Bigdely-Shamlo et al., 2015; Andersen, 2018; Jas et al., 2018). With the exception of bad channel removal and EOG selection, which was performed by visual inspection, the EEG pipeline relies on automated steps in order to improve both sharing of the results and further replication. We removed 13 peripheral electrodes (F11, FT11, T9, TP11, P09, I1, Iz, I2, P010, TP12, T10, FT12, F12), out of the 115 EEG electrodes, due to global bad impedance quality or no contact with the scalp at all. We low-pass filtered the raw data at 40 Hz cutoff frequency using a Finite Inpluse Response (FIR) filter and referenced to a common average. On the remaining 102 channels, sensors showing no signal, constant white noise or intermittent signal, as observed during the EEG recording, were interpolated. Cue onset was adjusted for the delay (+15 ms) between trigger generation and image appearance on the screen in the Faraday’s cage, measured using photodiodes during preliminary tests. We then segmented the trials from −1000 ms to 3500 ms around the cue onset. We systematically rejected trials with a large threshold of 350 *µ*V before detecting and correcting artifacts using the Autoreject library (Jas et al., 2017) with local thresholds detection and the Bayesian optimization method. Finally, we applied ICA correction to reduce the remaining eye blink artifacts based on one manually selected EOG channel.

### 4.11 Time-frequency

Our analysis focused on the contrast between No-Think and Think conditions during the TNT task for items classified as disgusting. For the ‘Think’ condition, we selected only trials where the participant reported a successful memory reactivation (feedback = 1). We kept all available ‘No-Think’ trials and extracted the power for each epoch in the high theta frequency band [5-9 Hz]. To do so we applied 5 Morlet wavelets in frequencies ranging from 5 to 9 Hz with a width of 5 cycles. After this procedure, we cropped the epochs from −0.5 s to 3.0 s to avoid edge artifacts and baseline corrected by subtracting the log ratio of the response in the interval from −500 to 0 ms. The resulting time-frequency data were then down-sampled to 10 ms time-bins for further analysis.

### 4.12 Statistical analysis

Analyses were performed using custom Python scripts (Python 3.6) together with specific modules for EEG analyses (Gramfort, 2013) and R packages (R version 3.4) for factorial analyses (Singmann et al., 2016) and effect size calculation (Torchiano, 2017). Robust correlations were performed using the correlation toolbox (Pernet et al., 2013) implemented in Matlab 2017b (The Mathworks). This method allows to reject the null hypothesis based on the percentile bootstrap CI, an approach less sensitive to the heteroscedasticity of the data than the traditional t-test. Figures, scripts, statistical analyses and part of the data reported in this paper are made available at: https://github.com/legrandnico/cardiaccontrol.

### 4.13 Behavior

Items that were not correctly recalled during the final test before TNT were systematically excluded from later analyses (Study 1: 1.2%; Study 2: 1.1%). We analyzed behavioral results with ANOVA integrating all the available factors and tested specific hypotheses with planned comparisons. We performed two-sided comparisons when no *a priori* hypothesis was formulated and one-tailed when we expected specific effects. In order to improve the clarity of our presentation, figures do not report all the possible comparisons but only relevant effects.

### 4.14 ECG

Comparisons between Neutral and Emotional heart rate curves were made using a paired t-test at each time point, we used the *fdr correction()* function implemented in MNE Python (Gramfort, 2013) to correct for multiple comparisons. For each group, we set the significance threshold at 0.05 to assess the effect of emotions in the first session and used a one-tailed t-test for measuring the effect of suppression in session 2.

### 4.15 EEG

We subtracted the mean values observed at each time point and for each electrode under the ‘Think’ condition to the ‘No-Think’ condition. We then performed a non-parametric spatiotemporal cluster statistic (Maris and Oostenveld, 2007) using implementations into MNE python as described in (Jas et al., 2018). We used a two-tailed t-test as the statistic on which to perform the clustering with permutation (n = 1024). As multiple clusters were outputted as significant, we automatically selected the cluster with the longest time duration.

## 5 Acknowledgments

We thank Lea Lemoine for his assistance with data preprocessing in Study 1, Emmanuelle Duprey and Camille Rebillard for the medical examinations and Florence Fraisse for administrative support.

## 7 Author contributions

N.L. programmed the experiment, collected and analyzed the data, and wrote the paper.P.G. designed the experiment, supervised data analysis, and wrote the paper. O.E. designed the experiment, supervised data collection, and provided inputs for data analysis and manuscript redaction. M.P., A.V., and F.V. performed medical inclusion visit. P.C. and F.D. supervised and performed EEG data collection. D.P. and F.E. supervised N.L.

### A Behavioral results - Study 2

We applied the same sequence of analyses for the behavioral data in Study 1 and Study 2.

#### A.1 Intrusions

We tested if the item categorized as disgusting produced less intrusion than neutral scenes as we previously observed. We compared the frequency of intrusion of Disgust and Neutral items over the eight TNT blocks for the No-Think condition (see Figure 1 - A in Appendix A.3). An Emotion x TNT blocks ANOVA showed both an effect of Emotion (F_(1, 23)_ = 19.49, 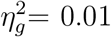, *p <* 0.001) and TNT blocks (F_(3.82, 87.88)_ = 24.75, 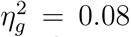, p *<*0.001) but no interaction between these two factors (F_(4.81, 110.61)_ = 0.73, 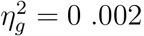, p = 0.60).

#### A.2 Inhibition of recall induces forgetting

An Emotion × Condition ANOVA on participants’ memories reflected by the Yes/No answer (see Figure 1 - B in Appendix A.3) revealed a significant effect for Emotion (F_(1,23)_ = 12,17, 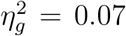, *p* = 0.002), Condition (F_(1.56,35.89)_ = 16.85, 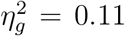, *p <* 0.001) but no interaction between those two factors (F_(1.44,33.05)_ = 2.52, 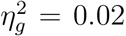, p = 0.11). Comparisons indicated that participants recalled significantly less No-Think item than Baseline for Disgust (t_(23)_ = −4.40, *p* = 0.0001, *d* = 0.89, one-tailed), while this effect was not significant with neutral items (t_(23)_ = −1.59, *p* = 0.06, *d* = 0.32, one-tailed). This result parallels previous works (Depue et al., 2007; Lambert et al., 2010) suggesting that emotional stimuli can be efficiently controlled and forgotten. These patterns of behavioral findings replicate observations made in Study 1 and suggest that emotional disgust may increase the motivation and the desire to reduce the momentary awareness associated with this unpleasant and unwanted memories. However, similarly to results reported in Study 1, disgusting items were globally more forgotten than neutral ones, which suggests that both a diminished memory of the items and a stronger motivation to control them could have produced the observed patterns.

#### A.3 Effect of suppression in the subjective evaluation

We then examined the SAM valence rating after the Think/No-Think procedure adjusted by pre-TNT evaluations (see Figure 1 - C in Appendix A.3). An Emotion × Condition ANOVA revealed a significant effect of Emotion (F_(1, 23)_ = 9.78, 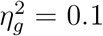, *p* = 0.005) but not for Condition (F_(1.97, 45.38)_ = 1.43, 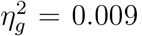, *p* = 0.25) or the interaction between those two factors (F_(1.94, 44.63)_ = 1.42, 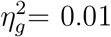, *p* = 0.25). Planned comparison showed a significant higher valence rating for No-Think items as compared to Baseline items for Disgust (t_(23)_ = 1.8, *p* = 0.04, *d* = 0.36, one-tailed). A difference was also observed for this emotion between Think, and Baseline images (t_(23)_ = 1.66, *p* = 0.05, *d* = 0.33, one-tailed) while no difference was found between Think and No-Think conditions (t_(23)_ = 0.01, *p* = 0.98, *d <* 0.01, two-tailed). Concerning Neutral scenes, we did not found similar increase of the valence rating for No-Think items as compared to baseline (t_(23)_ = 0.74, *p* = 0.46, *d* = 0.15). These findings show that suppressing disgusting scenes from memory may also alter the emotional quality of those memories so that their reappearance triggers less negative affect. This result parallels a recent study showing affective devaluation of words and objects after a TNT procedure (Vito and Fenske, 2017). Interestingly changes in affect do not arise for Neutral scenes suggesting that suppressing unpleasant memories may entail additional inhibitory effects not present for Neutral memories. Retrieval during Think trials measurably alters the perceived affect of the scenes to some extent, suggesting the existence of some form of habituation effect following repeated retrieval.

### B Behavioral indices of inhibitory control and evolution of emotional appreciation in Study 2 (n = 24)

### C Study 1 - NEUTRAL NAPS reference

Animals [ 043 h, 088 h, 097 h, 105 h, 108 h, 132 h, 155 v, 213 h], Faces [ 060 h, 061 h, 065 h, 123 h, 184 h, 211 h, 219 v, 292 h, 306 v], Objects [ 029 h, 049 h, 164 h, 200 h, 242 h, 270 h], People [ 026 h, 028 h, 035 h, 091 h, 095 h, 097 h, 100 h, 148 h, 149 h, 151 h, 164 h, 179 h, 249 h].

### D Study 1 - DISGUST NAPS reference

Animals [ 039 h, 041 h, 062 h, 065 h], Faces [ 156 h], Objects [ 006 h, 010 h, 011 h, 022 h, 125 h, 126 h], People [ 019 v, 077 v, 090 v, 205 v, 209 v, 220 h, 239 h].

### E Study 1 - SADNESS NAPS reference

Animals [ 013 h, 053 h, 067 h, 077 h], Faces [ 011 h, 012 v, 013 h, 019 h, 034 h, 149 v, 279 h, 294 h], People [ 001 h, 002 v, 127 h, 133 h, 143 h, 147 h].

### F Study 2 - NEUTRAL NAPS reference

Animals [ 043 h, 088 h, 097 h, 105 h, 108 h, 132 h, 155 v, 213 h], Faces [ 060 h, 061 h, 065 h, 123 h, 184 h, 211 h, 219 v, 292 h, 306 v], Objects [ 029 h, 049 h, 164 h, 200 h, 242 h, 270 h], People [ 026 h, 028 h, 035 h, 091 h, 095 h, 097 h, 100 h, 148 h, 149 h, 151 h, 164 h, 179 h, 249 h].

### G Study 2 - DISGUST NAPS reference

Animals [ 008 v, 018 h, 039 h, 041 h, 062 h, 065 h, 078 h], Faces [ 156 h, 296 h, 368 h], Objects [ 006 h, 007 h, 010 h, 011 h, 013 h, 021 v, 022 h, 023 v, 088 h, 106 v, 125 h, 126 h, 154 h], People [ 019 v, 077 v, 090 v, 204 v, 205 v, 209 v, 213 v, 217 h, 220 h, 223 h, 230 h, 239 h, 247 v].

